# Diagnostic Ion Data Analysis Reduction (DIDAR) allows rapid quality control analysis and filtering of multiplexed single cell proteomics data

**DOI:** 10.1101/2022.02.22.481489

**Authors:** Conor Jenkins, Benjamin C. Orsburn

## Abstract

Recent advances in the sensitivity and speed of mass spectrometers utilized for proteomics and metabolomics workflows has led to a dramatic increase in data file size and density. For a field already challenged by data complexity due to a dependence on desktop PC architecture and the Windows operating systems, further compromises appear inevitable as data density scales. As one method to reduce data complexity, we present herein a light-weight python script that can rapidly filter and provide analysis metrics from tandem mass spectra based on the presence and number of diagnostic fragment ions determined by the end user. Diagnostic Ion Data Analysis Reduction (DIDAR) can be applied to any mass spectrometry dataset to create smaller output files containing only spectra likely to contain post-translational modifications or chemical labels of interest. In this study we describe the application DIDAR within the context of multiplexed single cell proteomics workflows. When applied in this manner using reporter fragment ions as diagnostic signatures, DIDAR can provide quality control metrics based on the presence of reporter ions derived from single human cells and simplified output files for search engine analysis. The simple output metric text files can be used to rapidly flag entire LCMS runs with technical issues and remove them from downstream analysis based on end user minimum requirements. Acquisition files that pass these criteria are further improved through the automatic removal of spectra where insufficient signal from single cells is observed. We describe the application of DIDAR to two recently described multiplexed single cell proteomics datasets.

**Abstract Graphic:** 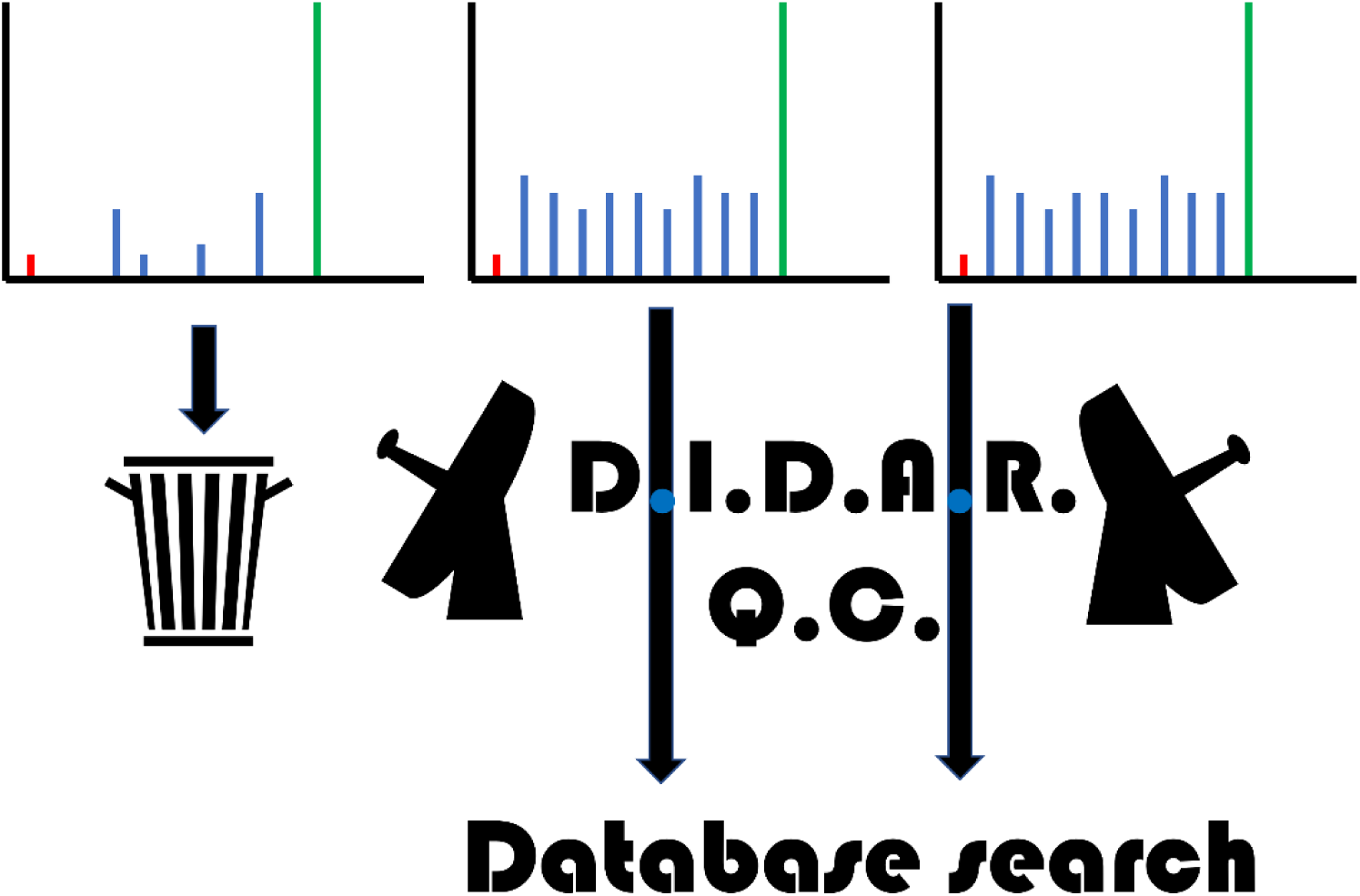

## Introduction

Liquid chromatography coupled to high resolution tandem mass spectrometry (LCMS) continues to remain the dominant technology in proteomics^2^, metabolomics^3–5^ and lipidomics^6^ research. Nearly all proteomics and metabolomics data processing today is performed using desktop personal computer (PC) architecture.^1^ While many open software packages are amenable to installation on high performance cluster environments (HPCs)^7–9^ these have yet to be adopted by the LCMS community. Likewise, software designed for use on the Cloud have been demonstrated but have been slow to be adopted despite the promise of improved speed and increased biological relevance processing on the Cloud can allow.^10,11^ One method to circumvent the size of the generated LCMS files is to apply pre-filters to remove tandem mass spectra (MS2) that are of interest. We have recently described Reporter Ion Data Analysis Reduction (RIDAR) which allows us to set minimum quality filters and relative quantification values in isobarically tagged proteomic datasets. Spectra lacking reporter ions of sufficient signal to noise ratio across a user defined minimum number of reporters can be discarded. In addition, spectra that are not quantitatively interesting to the end user, such as peptides that display no change in relative quantities can likewise be filtered prior to data analysis.^12^

Diagnostic oxonium ions produced during the fragmentation of glycopeptides are commonly used in both real time data acquisition and in data processing. Orbitrap systems can identify ions producing oxonium ion fragments for further analysis, with the most impressive results obtained by further analysis with electron transfer dissociation techniques.^13,14^ Software such as SugarQB begin by separating out MS2 spectra with apparent oxonium ions for specialized searches considering the glycan moieties that could lead to the production of those ions.^15^ Although not as frequently used, many other commonly studied post-translational modification also produce unique diagnostic fragment ions that could be utilized in a similar manner. A systematic analysis of modified peptides synthesized for the ProteomeTools project has identified fragment characteristics and diagnostic ions for many PTMs.^16^ The utilization of diagnostic ions can help prevent the misidentification of modified peptides such as arginine citrullination, which has been the topic of multiple controversial studies and recent retractions.^17–19^

Single cell proteomics by mass spectrometry is a new field of research largely made possible innovative applications of existing technology. The revolutionary work of Budnik *et al*., first described the SCoPE-MS workflow whereby the additive signal of multiplexed peptide samples could be leveraged to effectively amplify peptide signal.^20^ To achieve these effects SCoPE-MS employs a carrier channel of isobarically labeled peptides of higher relative concentration to resuspend labeled peptides from single cells. Data dependent LCMS analysis of these multiplexed samples leads to the fragmentation of and identification of peptides from the carrier channel. The isobaric reporter ion fragments corresponding to labels applied to single cells, if within the intrascan linear dynamic range of the instrument, can be used for both peptide identification and quantification.^21–24^ Although other single cell technologies such as single cell RNAseq have been sorting and analyzing single cells for over a decade with increasingly sophisticated technology, relatively little of this can be applied to proteomics approaches.^25,26^ While new technologies such as nanoPOTs^27^, nPOP^23^ and ProteoChip^24^ are improving these aspects, technical challenges in sample preparation may be the biggest limiter to entry into single cell proteomics. Single human cells and the volumes of reagents used in their preparation and analysis can be affected by factors as ubiquitous as relative humidity and static electricity. As a relatively large number of single cells must be analyzed for sufficient statistics to capture both technical and biological variability, failed runs may not be detected until data analysis is completed which may take days or weeks on desktop PC architecture. Quality control tools that could be leveraged prior to complete data analysis could simplify and improve nearly all downstream processes.

## Methods and Materials

### Script development

DIDAR was written in Python 3.8 using the Spyder 4.1.4. IDE.^28^ DIDAR is contained in less than 100 total lines of code inclusive of comments and line spacing. In addition, utilizing the built-in os python package as the only dependency adds to the lightweight nature of the program.

### Single cell proteomics by mass spectrometry (SCoPE-MS) data quality filtering

For the development of DIDAR as a quality control analysis pipeline for SCoPE-MS we used publicly available files from two recent SCoPE-MS studies. The first utilized the SCOPE2 methodology using both TMT11-plex and TMTPro16-plex tags to study macrophage heterogeneity using a quadrupole Orbitrap MS2 based approach and 95 minute acquisition times. The second used the same sample preparation method but utilized the 9 unit resolution mass tags from the TMTPro16 reagents to study single human KRASG12C model cells (NCI-H-358) using a third-generation TIMSTOF mass analyzer and 30 minute acquisition methods. The vendor format instrument files from each study were converted to MGF using MSConvert Version: 3.0.20310-0d96039e2 (developer build) using the default conversions for each instrument. For applying DIDAR to these SCoPE-MS files, the DIDAR.py and diagnostic ion text was copied into the folder containing the MGF files from each respective study. The diagnostic ion text file referenced by DIDAR only contained the exact masses of the reporter ions as reported by the manufacturer. The DIDAR script was executed identically for each file, with the exception of the ion mass tolerance which was set at 0.002 Da for the Orbitrap files and 0.005 for the TIMSTOF files to compensate for the relative differences in mass accuracy between these analyzers.

### Data Availability Statement

All original files used in this study are publicly available on the MASSIVE repository (massive.ucsd.edu). The SCOPE2 quadrupole Orbitrap files are available as accessions MSV000083945 and MSV000084660. The TIMSTOF single cell files were accessed as accessions MSV000088144 using the reviewer password provided in the study preprint (pasefSCOPE700N). All DIDAR filtered MGFs and output metrics for this study have been deposited at: ftp://massive.ucsd.edu/MSV000088887/.

### Code availability statement

The DIDAR script and diagnostic ion text files for TMT reporter ions and video tutorials are available at: https://github.com/orsburn/DIDARSCPQC.

## Results and Discussions

### Reanalysis of two SCoPE-MS studies

Twenty five LCMS files from two separate SCoPE-MS based studies were obtained from public repositories and analyzed with DIDAR using the corresponding exact masses of the reporter ions used in the studies as the filtering parameters. A summary of the numbers of spectra exhibiting each reporter ion across these twenty five files is shown in **Figure 1A/B** and full summary DIDAR results are available as **Supplemental Data 1**.

**Figure 1.**
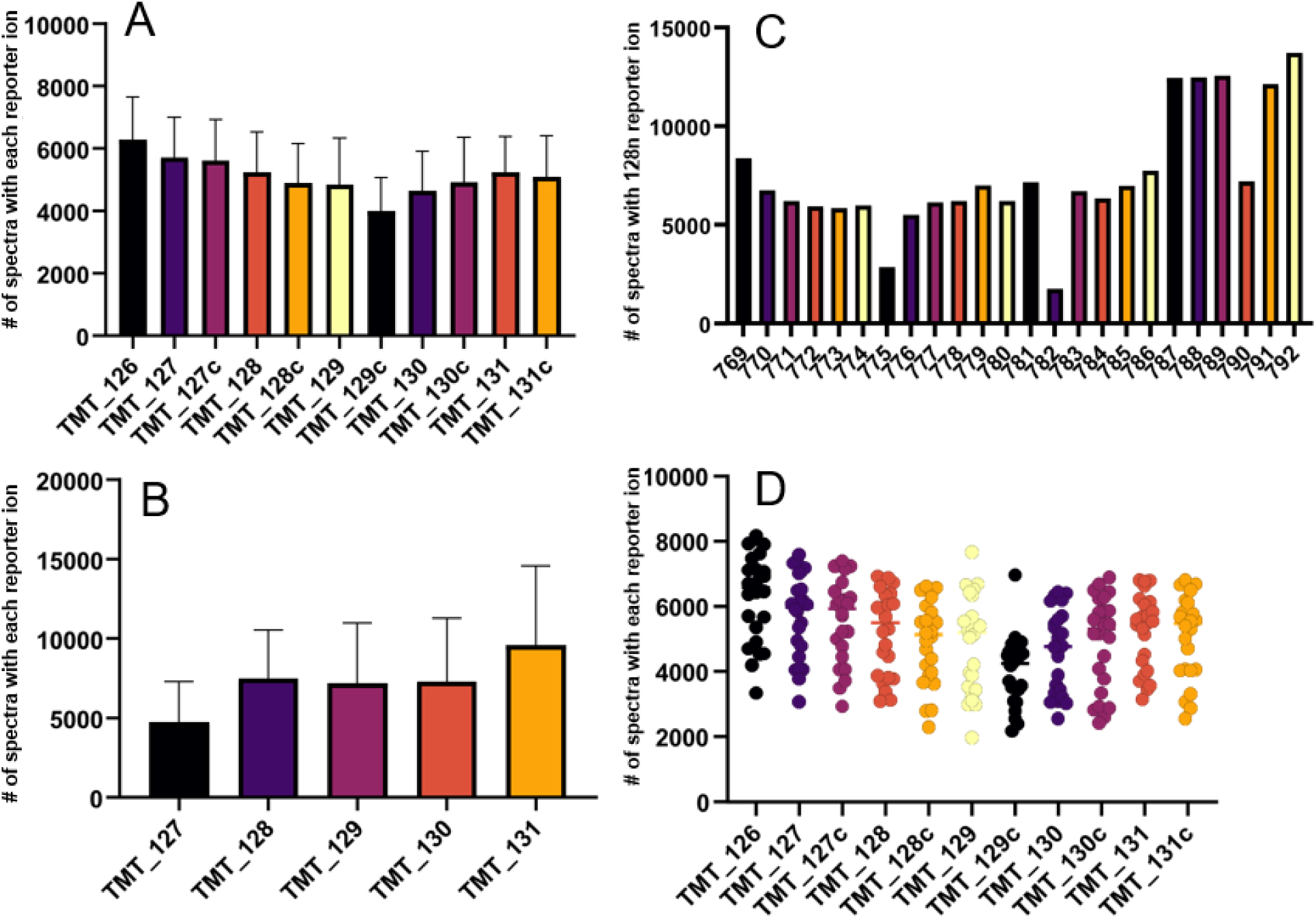
A comparison of DIDAR diagnostic results from two recent multiplexed single cell proteomics studies. **A**. A study that utilized a Mantis liquid handler for sample prep and Orbitrap based analysis using 100 minute instrument acquisition time. **B**. A study where single cells were prepared by hand in microliter volumes and analyzed on 30 minute gradients with a time of flight analyzer. **C**. An illustration of the use of a single reporter ion count as a QC metric. **D**. A plot demonstrating the variability across a batch of over 250 single cells.

While both approaches used the same basic sample preparation procedure, one study used an automated liquid handler for sample prep while the second lysed and labeled cells by hand. In addition, the two studies used very different chromatography and mass spectrometry, with the earlier SCOPE2 study employing quadrupole Orbitrap technology and approximately 100 minute methods. The second study used a methods a total of 45 minutes in length and a TIMSTOF for 9-plex reporter ion quantification. As shown in **Figure 1A/B**, the number of spectra with each reporter ion appears to be more consistent in the original SCOPE2 study compared to the more recent pasefRiQ analysis.

### DIDAR can flag outlier channels and provide metrics of data reliability

Each file processed through DIDAR is assigned two output files. The first is a new filtered MGF containing only spectra meeting the end user’s criteria. For example, by changing the minimum number of reporter ions to 5, we will create an output file that only contains spectra were at least 5 of the TMT reporter ions are observed. In this study we left the default number of ions at 1 to capture the widest variability in overall metrics. The second output file is a short diagnostic text file that provides a simple count of the number of spectra where each diagnostic ion is counted. **Figure 1C** is a visualization form the 25 pasefRiQ files analyzed. As shown, both files 775 and 782 have a number of 128n reporter ions that appear significantly below the average of the other files analyzed within this batch. In addition, we can observe a general increase in the number of spectra with reporter ions at later points in this batch. General trends of increasing or decreasing numbers of reporter ion signal may indicate systematic errors in sample handline or other hardware. When evaluating the original diagnostic counts output data (**Supplemental Data**) we see that in run 782 the failure was comprehensive for the entire experiment as no cells returned more than 1,500 spectra with observed reporter ion signal. In contrast, run 775 appears to have a near average number of reporter ions in all measured channels except 128n, suggesting a technical issue occurred in the handling or processing of that single cell. DIDAR output can also be combined to evaluate the reproducibility of data within and between batches. As shown **Figure 1D**, these data can be easily examined for simple analyses of each individual cell and file across batches of hundreds of individual cells.

## Conclusions

We have described the application of a rapid lightweight Python tool for the quality assessment and filtering of single cell proteomics data. When applied in this manner, DIDAR can flag both entire LCMS runs that experienced technical issues for further analysis or for removal from analysis. In addition, DIDAR can generate new input files that only contain data meeting user required minimum cutoffs.

